# Probing Protein Allostery as a Residue-specific Concept via Residue Perturbation Maps

**DOI:** 10.1101/355370

**Authors:** Hamed S Hayatshahi, Emilio Ahuactzin, Peng Tao, Shouyi Wang, Jin Liu

## Abstract

Allosteric regulation is a well-established phenomenon classically defined as conformational or dynamical change of a small number of allosteric residues of the protein upon allosteric effector binding at a distance. Here, we developed a novel approach to delineate allosteric effects in proteins. In this approach, we applied robust machine learning methods, including Deep Neural Network and Random Forest, on extensive molecular dynamics (MD) simulations to distinguish otherwise similar allosteric states of proteins. Using PDZ3 domain of PDS-95 as a model protein, we demonstrated that the allosteric effects could be represented as residue-specific properties through two-dimensional property-residue maps, which we refer as “residue perturbation maps”. These maps were constructed through two machine learning methods and could accurately describe how different properties of various residues are affected upon allosteric perturbation on protein. Based on the “residue perturbation maps”, we propose allostery as a residue-specific concept, suggesting all residues could be considered as allosteric residues because each residue “senses” the allosteric events through perturbation of its one or multiple attributes in a quantitatively unique way. The “residue perturbation maps” could be used to fingerprint a protein based on the unique patterns of residue perturbations upon binding events, providing a novel way to systematically describe the protein allosteric effects of each residue upon perturbation.

**Author Summary:** Allostery is protein regulation at distance. A perturbation at one site of the protein could distantly affect another site. The residues involved in these sites are considered as allosteric residues. The allostery concept has been widely used to understand protein mechanisms and to design allosteric drugs. It is long believed only a small number of residues are allosteric residues. Here, we argue that all residues in a protein are allosteric residues. Upon the perturbation of the allosteric events, the different properties of each residue are affected at the distinct extend. We used hybrid models including molecular dynamics simulations and machine learning components to reveal that not only many properties of residues are affected upon ligand binding, but also each residue is affected through perturbation of its various properties, which makes the residue distinguishable from other residues. According to our findings in a model protein, we defined a “residue perturbation map” as a two-dimensional map that fingerprint a protein based on the extent of perturbation in different properties of all its residues in a quantitative fashion. This “residue perturbation map” provides a novel way to systematically describe the protein allosteric effects of each residue upon perturbation.

## Introduction

Allostery has been an evolving concept (1) in proteins from considering the allosteric proteins as two-state switches that were either conceitedly (2), or sequentially (3) affected by an effector molecule, to considering them as conformational ensembles whose populations are shifted upon binding to an effector (4–6). More modern views propose that all proteins are intrinsically allosteric (7) as subtle conformational changes can occur upon binding to a ligand in a classically-considered non-allosteric protein. From a mechanistic view point, it can be speculated that binding to an effector propagates a signal through changing properties of a network of receptor residues, including their conformational dynamics (8) and nonbonding interaction attributes (9), which may not even accompany observable conformational changes (10). Based on these insights, it is reasonable to hypothesize that the binding of a ligand to a protein changes multiple attributes of a network of so-called “allosteric” residues. Such broader definition, if shown to be true, would raise the question that whether all such different attributes are perturbed in all affected residues, or alternatively, each residue or sets of residues are affected in a different way. The latter possibility can be described in other words as different allosteric residues “sense” effector binding to a pocket in different ways. With this broader view, it can also be further hypothesized that all residues in a given protein are potentially allosteric, although the quality and extent of their communication with the binding site varies.

Here, we tested these hypotheses in the PDZ3 domain of PSD-95, which is a well-known model for analyzing allosteric effects (9, 11–18). Two crystal structures of PDZ3 in bound and unbound states are available (19). The bound structure includes a five-residues peptide ligand, which is bound in a groove walled with a helix (αB) and a sheet (βB). Petit *et al.* have used Isothermal Titration Calorimetry (ITC) and Nuclear Magnetic Resonance (NMR) to highlight some dynamic allostery relative to an α-3 helix in PDZ3 (12). Gerek and Ozkan have highlighted some residues involved in allosteric pathways using Perturbation Response Scanning (PRS) (14). McLaughlin *et al.* have identified different mutational routes that could affect the binding specificity in PDZ3 via a high-throughput single mutation analysis (15). In another research work, Murciano-Calles *et al.* reported the allosteric effects of post-translational modifications of some residues in PDZ3 (17). Recently, computational methods have also been employed by different research groups to study the allosteric effects in PDZ3. Among these works is a Monte Carlo path generation simulation by Kaya *et al.* which has revealed some potential propagation routes of the allosteric signals in some proteins including PDZ3 (16). Kalescky *et al.* have used Molecular Dynamics (MD) simulations to perform a rigid residue scan in the bound and unbound PDZ3 and identified some allosteric residues that matched with previous experimental observations (18). A more recent study by Kumawat and Chakrabarty has reported the use of MD simulations and consequent revealing of the electrostatic interactions as the most significant hidden basis of dynamic allostery in PDZ3 (9).

With the question about the attribute/residue-specific perturbations upon ligand binding, we propose the use of hybrid models that could estimate the extent of property perturbations in different residues upon ligand binding. One set of such models use deep learning neural networks to compare the possibility of prediction of binding status in PDZ3 with different residue properties as descriptors. These models reveal whether different residue attributes such as position, nonbonding interaction and dynamics at various time scales have the same or different ability to predict the binding status, and hence are affected upon binding similarly or in different levels. The other set of models use the variable selection capability of random forests to rank the residues described with different attributes in terms of their contribution to distinguishing the protein binding state. These models assign significance indices to each residue in terms of its different properties and show whether different attributes are perturbed to the same extent in all affected residues. We present the significance indices as two-dimensional diagrams, which we name Residue Perturbation Maps. These maps fingerprint each protein in terms of its interaction with a specific binding ligand in a visual easy-to-follow fashion.

What empowers the machine learning models in our approach is that we integrate them with MD simulations, which provide a sufficient sampling pool of snapshots as training and testing records to train and validate deep neural networks and random forest models. Machine learning models have been used widely to study the structures and interactions among biomolecules (20, 21). However, the challenge in using these methods, especially the deep learning neural networks is the availability of thousands of samples needed to train them. This study is proposed to investigate the possibility of using snapshots from bound and unbound MD trajectories to calculate several residue attributes, which then could apply machine learning models to test the mentioned hypotheses. The scheme of this approach is represented in Figure 1.

**Figure 1.**
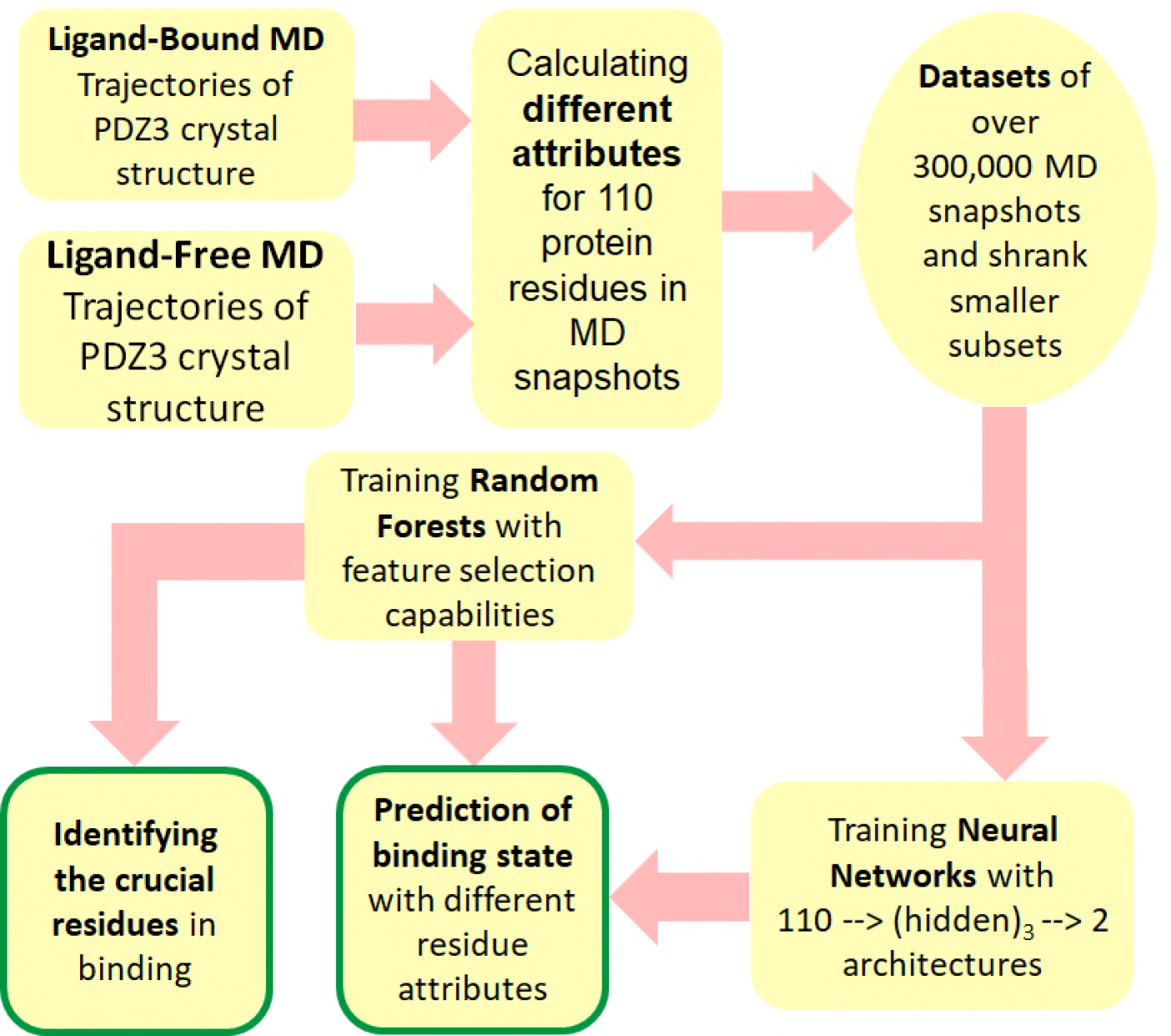
Scheme of the approach proposed in this work.

## Methods

### MD simulations and analysis

Residues 306-415 in the crystal structures of the bound and unbound PDZ3 (PDB codes: 1BE9 and 1BFE, respectively) were used as initial structures, and the rest amino acids were removed. The program cpptraj (22) of Ambertools 16 (23) was used to analyze the trajectories. Details about MD simulations and analysis are presented in the supporting information.

Different types of descriptors defined as follows were generated using combinations of cpptraj and python scripts for 110 residues (#306-415) of each simulation snapshot. To describe the position, the distance between the C_α_ atoms (C_α_ model) as well as the distance of the residues’ geometric centers (GEOM model) and the protein geometric center were calculated. The former would describe the changes in the backbone position, while the latter would incorporate the side chain positions in the calculations. The residue fluctuations (FLUCT model) were measured as the root mean square distance (RMSD) between the residue heavy atoms at each snapshot and its averaged structure in a single trajectory. The non-bonding properties were defined as the van der-Waals (VDW model), electrostatic (ES model) and their combined interactions (NONB model) between each residue and the rest of the free protein in each snapshot. The residue surface area (SURF model) was described as the contribution of the surface area of each amino acid to the surface area of the free protein (ligands were ignored in ligand-bound snapshots). The residue dynamics (DYN models) were measured as running RMSD values over the MD trajectories with 10 ps, 1 ns, 25 ns and 100 ns window sizes. Table 1 summarizes these descriptors and their abbreviations as used throughout the article. Each model was saved as a CSV file for all 338538 snapshots. Sample trajectory analysis scripts are provided in the supporting information.

### Deep Neural networks and random forests

Neural networks were trained and tested in 10 iterations of 5-fold cross validations in which the records were shuffled in each iteration and 80% were used for training while 20% were used for testing in each cross-validation iteration. To perform random forest modeling, we used the random forest classifier from the Scikit-Learn machine learning library (24). Details about the neural network and random forest models are presented in the supporting information.

## Results

### Multiple attributes of allosteric residues are affected upon ligand binding

Several properties can be calculated for a protein from the molecular dynamics snapshots at the domain, residue, sub-residue and atomic levels. These properties describe the system in different ways and could be considered as different layers of information that potentially result in a comprehensive picture of the protein when combined. From a protein allostery viewpoint, it is interesting to investigate the fluctuations of such properties at the residue level of focus. We hypothesized that each one of such layers of information can differentiate between the snapshots taken from the ligand-bound and ligand-free MD trajectories with different efficiencies. In other words, multiple residue attributes are affected upon ligand binding to different extent. To test such hypothesis, we described the residues of the PDZ3 protein in 2 μs of ligand bound and 2 μs of ligand-free MD simulations, in terms of their positions, fluctuation, nonbonding interactions, surface area and dynamics. **Error! Reference source not found.** summarizes different residue descriptors which were calculated from MD trajectories and used to train and test predictive machine learning models. More details are provided in the Methods section.

**Table 1.** Description of the properties (attributes) of residues in PDZ3 domain that were calculated from MD trajectories and used to train predictive machine learning models.

| Abbreviation | Description |
| --- | --- |
| $C_{\alpha}$ | Distance between $C_{\alpha}$ of the residue and the geometric center of the protein |
| GEOM | Distance between geometric center of the residue and the geometric center of the protein |
| FLUCT | Root mean square deviation (RMSD) between the residue and its average structure in the trajectory |
| VDW | van der Waals interaction between the residue and the rest of the protein |
| ES | Electrostatic interaction between the residue and the rest of the protein |
| NONB | Total non-bonding (VDW+ES) interactions between the residue and the rest of the protein |
| SURF | Contribution of the residue to the protein surface area |
| DYN-10 ps | Dynamics of the residue in 10 ps time scale |
| DYN-1 ns | Dynamics of the residue in 1 ns time scale |
| DYN-25 ns | Dynamics of the residue in 25 ns time scale |
| DYN-100 ns | Dynamics of the residue in 100 ns time scale |

We trained and optimized deep learning neural network models using the residue, which were descriptors generated and calculated from MD trajectories based on a 5-fold crossvalidation procedure that randomly separated the dataset into 80% for training and 20% for validation in each run and repeated five times. Snapshots taken every 10 ps in MD trajectories were used to train models with ability to distinguish the bound and unbound proteins in relatively high accuracies (Figure 2). To identify which residue attributes have more efficient distinguishing capabilities, we reduced the snapshots in the data sets by picking one snapshot in every 100 ps, 1 ns and 10 ns of MD trajectories (different snapshot offsets). The datasets containing maximum data points (saved with 10 ps intervals) were also used to train random forest models. The random forest models resulted in similar prediction accuracies as the neural networks (Figure 2). In terms of the residue descriptors, the residue fluctuations (FLUCT) resulted in the highest accuracy in all snapshot offsets. The surface area contributions (SURF) and the position descriptors (GEOM and C_α_) resulted in the second and the third highest prediction capabilities among others. Comparing the efficiency of models trained with GEOM and C_α_ implies that incorporating the information about the side chains (GEOM) slightly improves the prediction accuracy. The nonbonding interactions generally resulted in poor predictions comparing to other descriptors; however, the electrostatic interactions distinguished the binding states better than vdW interactions, implying that the electrostatic interactions are more affected upon ligand binding. The results showed that the efficiency of the dynamics properties depends heavily on the time scale of the dynamics being considered. Higher time scale dynamics resulted in much better prediction performance than low time scale dynamics.

**Figure 2.**
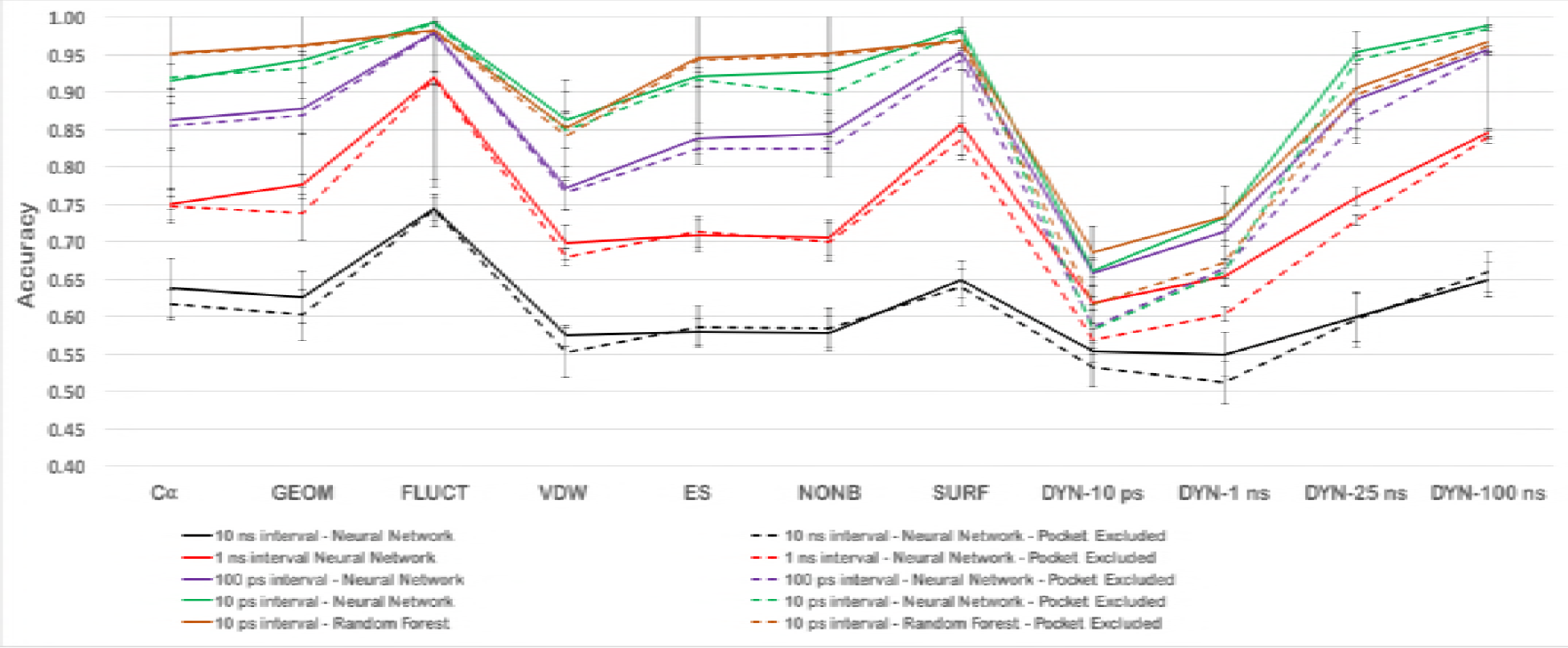
Accuracies of neural network and random forest models fed by different subsets of the MD snapshots and different residue descriptors. The reported neural network accuracies were averaged over ten average accuracies of cross validation attempts, each of which, the networks were trained with 80% of snapshots and tested with 20%. The random forest accuracies were averages of one hundred runs of independent random forest models. Error bars represent the standard deviation of independent runs.

It is possible that the differentiation capabilities mostly rely on information from the residues close to the binding site, rather than allosteric residues in farther distances. To understand the role of such distal residues in distinguishing the binding states, the same neural networks and random forests were trained without the data from the pocket residues. The pocket residues were defined as residues that are located within 3 Å distance from the bound peptide in the crystal structure (PDB code 1BE9). Such definition excludes 12 residues (323,324,325,326,327,328,339,372,373,376,379,380) from consideration for training the models. The accuracies of the predictive models did not drop dramatically when excluding the pocket residues (Figure 2). This shows that the ability of the neural networks to distinguish between the bound and unbound states does not mainly originate from the difference in residues near the binding site, rather, it relies on the allosteric resides. However, a higher pocket residue reliance of model accuracy can be observed when using the low timescale dynamics to train neural networks (Figure 2). The accuracies of all models that were trained with 10 ps and 1 ns dynamics data deteriorate significantly when the pocket residues are ignored. This is reasonable as the bound ligand is expected to change the low timescale fluctuations of residues that it directly interacts with.

### Allosteric residues are affected in different ways upon effector binding

Considering the inter-dependence between properties of protein residues and ligand binding events, it is expected that specific residues can potentially change the protein affinity for a ligand, and could be affected by the binding ligand at the same time. The residues with a better ability to “sense” ligand binding can be considered as potential allosteric residues. Various experimental and computational approaches have been used to identify such potential allosteric residues in model proteins such as PDZ3 (12–18). However, it is still interesting to know whether different allosteric residues sense the ligand binding in the same way (i.e. being affected through perturbation of the same residue properties), or they respond to perturbations through changes of different residue properties. Also from a more practical viewpoint, it is beneficial to know whether the changes in the same or different properties of these residues result in allosteric effects. Affected in different ways is schematically represented as in Figure 3.

**Figure 3.**
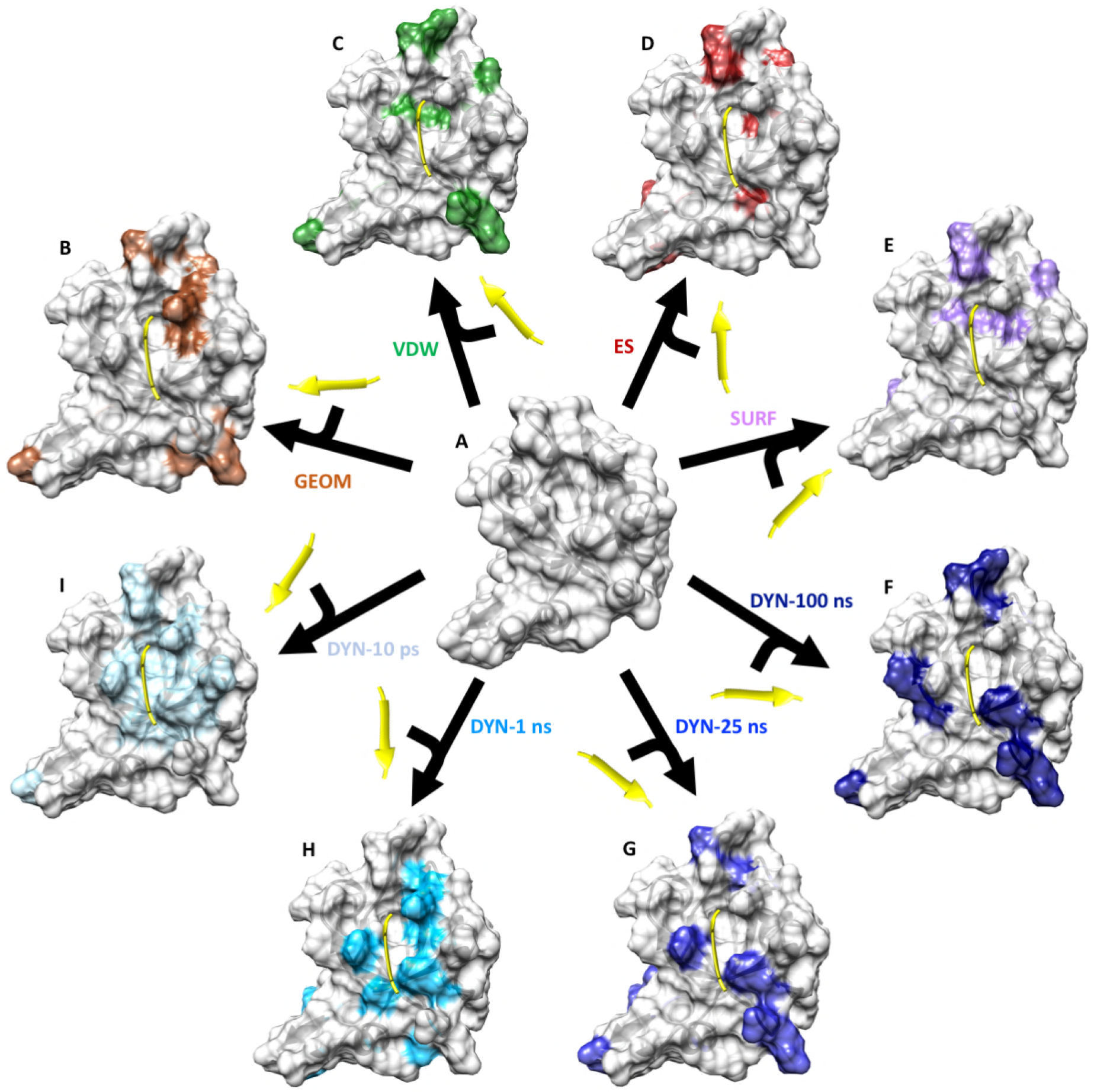
Schematic representation of how various residues are affected in different ways upon binding of the PDZ3 domain to its peptide ligand. The figure includes the crystal structures of the unbound protein (A), and the crystal structure of the bound protein (B-I) in different representations. As noted in the text, many random forest models were trained with different properties of the protein residues to distinguish the bound and unbound MD snapshots. The 15 most contributing residues selected by the random forest models are highlighted in each of the bound protein representations (B-I) with a distinguishing color.

To answer this question in the PDZ3 domain as a model protein, we used the random forests model with the inherent ability to quantitatively rank the contributions of residues for prediction purpose. Such contributions were extracted from the random forest models that were used to distinguish between the bound and unbound snapshots of MD trajectories. Figure 3 schematically represents the 15 most contributing residues in that were selected by some random forest models. We also represent these contribution values from random forest models quantitatively as color intensities in “residue perturbation maps” (Figure 4A, B). The results show that the affected residues have different contributions to model accuracies when different residue properties are used to train the models, which essentially indicates that not all protein residues are affected in the same way.

**Figure 4.**
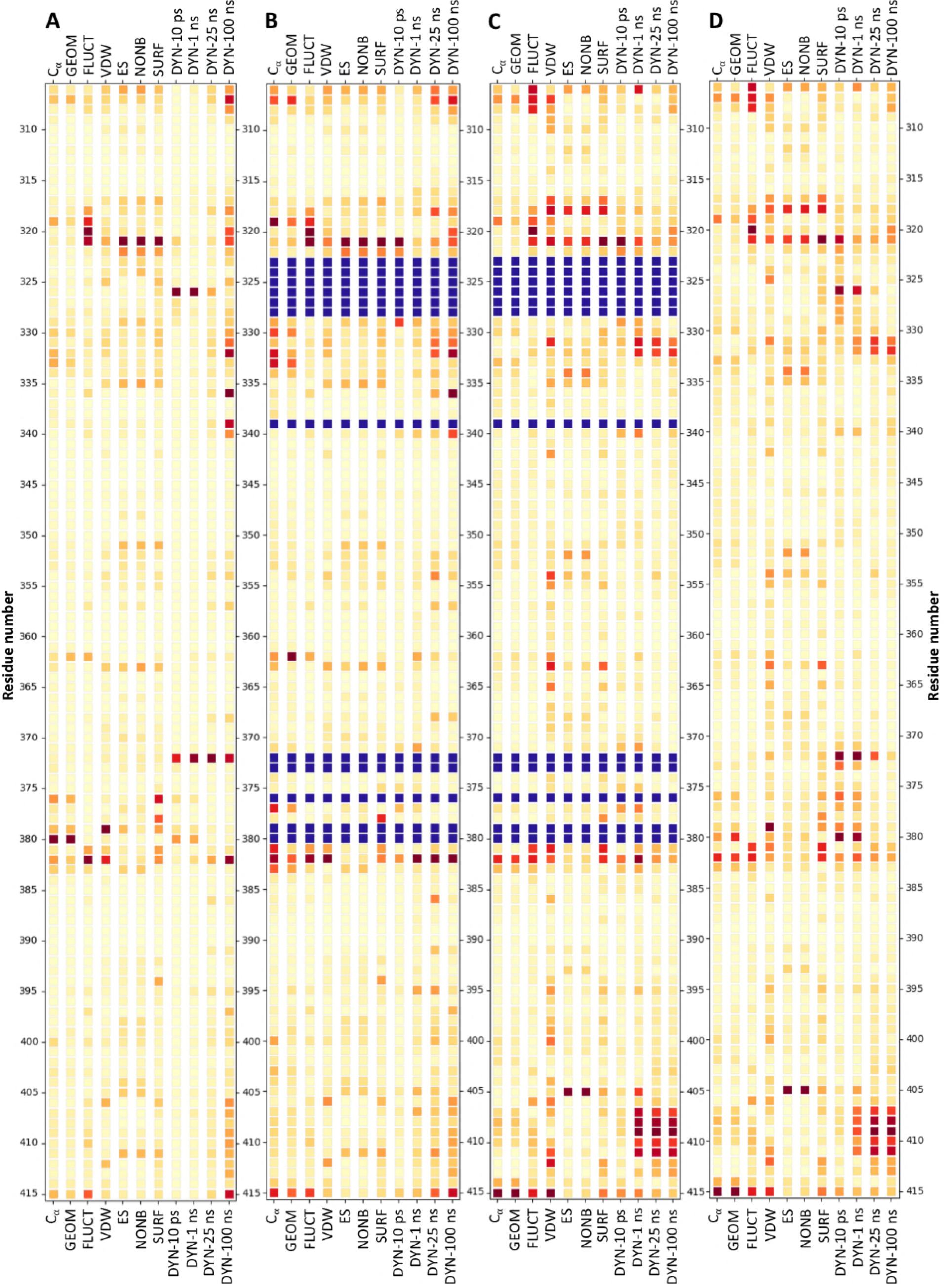
**A and B)** Residue perturbation maps representing the contribution of each residue in the accuracy of the random forest models with considering the pocket residues (A), and without considering the pocket residues (B). **C and D)** Residue perturbation maps calculated directly from MD trajectories represented as the absolute difference between the average property values in bound and unbound states, with considering the pocket residues (C) and without considering the pocket residues (D). Red intensity correlated with the contribution in accuracy in A and B and with absolute average differences in C and D. Residues in solid blue are considered as pocket.

### Residue Perturbation Maps with different flavors

Because of their occurrence at the ligand binding site, it is expected that the pocket residues undergo higher perturbations in terms of their attributes, relative to other residues, and hence they would contribute more to the random forest accuracies. This would mask the importance of other residues in allosteric positions in a perturbation map like the one in Figure 4A, because the data is normalized prior to representation in maps. Thus, we prepared the maps from random forest models that were trained both with and without the data from the pocket residues (Figure 4A vs. 4B, respectively). It is clear from the maps that ignoring the pocket residues highlights the importance of some other residues (darker color) that were pale when pocket residues were considered.

We also prepared the residue perturbation maps directly from the MD trajectories rather than indirectly from random forest model selected features to validate the random forest results. To do this, the absolute difference of the property values between bound and unbound trajectories were mapped again with and without considering the pocket residues (Figure 4C and 4D, respectively). Comparing the maps from raw MD trajectories and from random forest, importance values show that although many residue properties that are different in bound and unbound trajectories and were also identified as important factors in binding by random forest models, there is no one-to-one quantitative correlation between them. In fact, importance factors extracted from random forest models are mostly qualitatively highlighted in raw data, but random forests rank the residues in a different way relative to binding. This might be resulted from removing noises from the raw data with ignoring the differences not directly related to binding. For example, dynamics of the residues 407-411 which occur very close to the C-terminal of the protein and far from the binding site shows a significant difference between the bound and unbound trajectories. But these differences, especially at lower time scales, were not recognized as highly important in the classification of the bound and unbound states.

### Previously highlighted residues glow in Residue Perturbation Maps

Interestingly, many PDZ3 residues that were highlighted in the residue perturbation maps as top contributing ones in their predictive capabilities, or as just being different in MD simulations have been reported previously as allosteric residues. Residues 317-322 (Figure 5A,B) significantly contribute in the prediction of the binding status in many random forest models. Such high contribution is expected because the conformations of these residues are significantly different in the crystal structures (19), which were used as initial simulation structures. However, their contributions to prediction vary when using different descriptors. For example, the low timescale dynamics of Thr321 and Gly322 (DYN-10 ps) as well as their electrostatic (ES) and surface contributions (SURF) seem to be more important in prediction of binding state than their position GEOM and C_α_, while the position of Gly319 plays a more important role in prediction than its electrostatic interactions.

**Figure 5.**
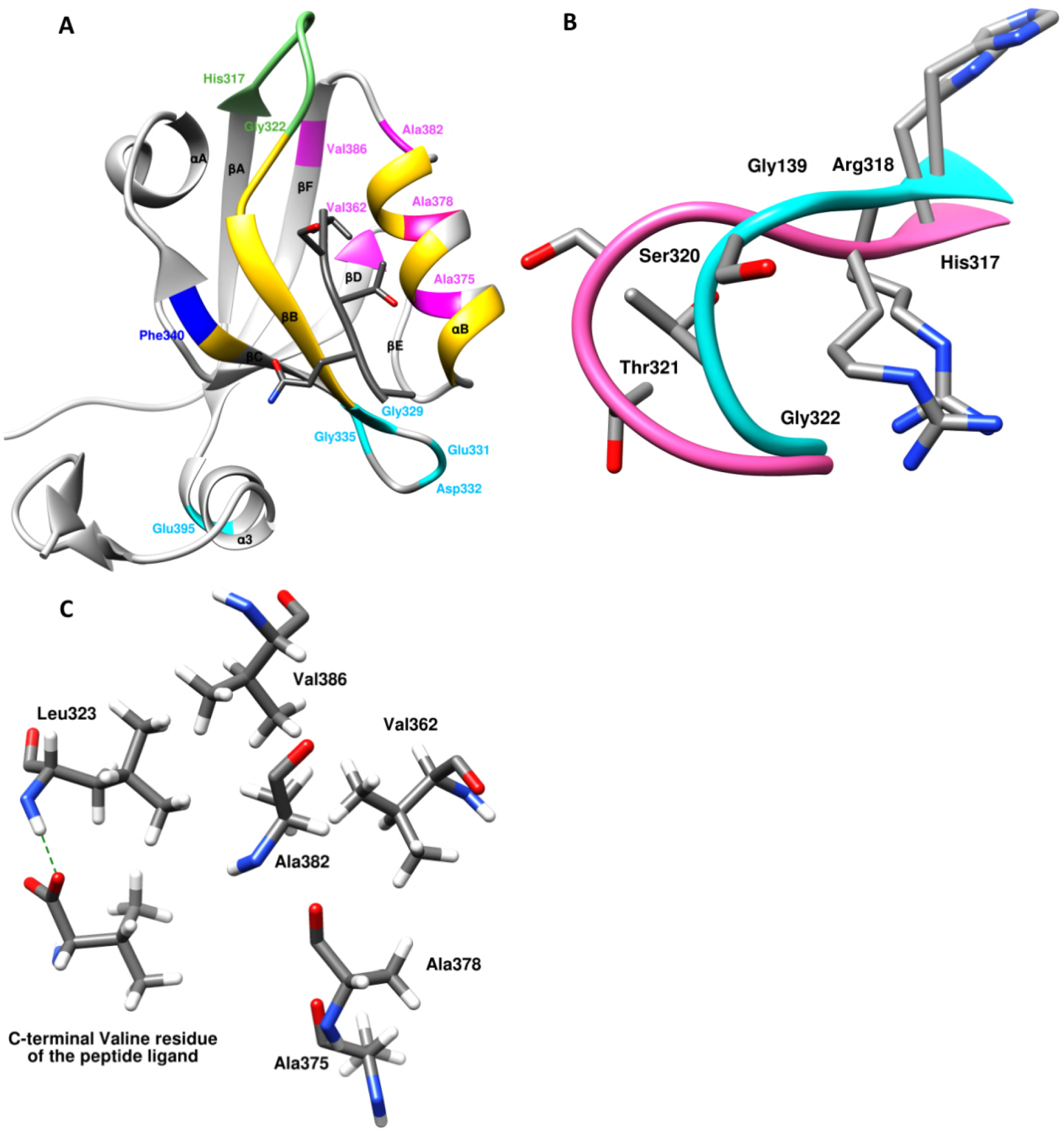
**A)** Ribbon representation of the crystal structure of PDZ3 protein (PDB code: 1BE9) highlighting some important residues recognized by random forest models. Residues in gold are considered as pocket residues and were not fed into the random forest models. B) Residues 317-322 in the ligand bound crystal structure (cyan) and unbound crystal structure (pink) of PDZ3, as the backbones of complete structures are superimposed. Hydrogen atoms are not shown. C) Localization of a network of hydrophobic residues of PDZ3 and the C-terminal of the peptide ligand. The green dashed line represents the key salt bridge between the peptide ligand and backbone of Leu323.

Gly329 has high importance in models using total nonbonding interaction (NONB) and low timescale dynamics (DYN-10 ps and DYN-1 ns) as predictive descriptors. This residue is reported to have the largest mutational effect in the protein (15) and relatively high allosteric response ratio as calculated by Grerek and Ozkan (14). Our finding implies that the Gly329 probably experiences different dynamics in bound and unbound states. This residue is located in the beta sheet (βB), which is involved in the formation of the binding pocket. So, it is reasonable to expect that its short-range nonbonding interactions and low time scale vibrations are affected by ligand binding.

It has been shown that phosphorylation of the residues at the α3 helix in PDZ3 (residues 394-399) play important roles in binding (12). Among the attributes of residues in this short helix, the dynamics of Glu395 at 1 ns time scale is shown to be affected by binding in our results. Gly335 which can interact with this helix only through nonbonding interactions, has been shown sensitive to its deletion in the ^15^N relaxation experiments (12). Our results suggest that Gly335 is affected by the ligand through nonbonding interactions (according to VDW, ES and NONB models). This fluctuation in nonbonding interactions might be related to the changes in low time scale dynamics of Glu395, as it is expected that the fluctuations of the negatively charged side chain of Glu395 affect its surrounding environment through nonbonding interactions. This can be considered as a hypothesis that could be evaluated in future research. In the crystal structure (19), Gly335 is close to Gly329 in space, which is in turn in close proximity to the ligand as mentioned above. This implies that the ligand affects low time scale vibrations and nonbonding interactions of Gly329, which affects Gly335 and the α3 residues in a sequential fashion through nonbonding interactions. This nonbonding effect on the α3 helix has not been recognized in our models, although some residues in the helix show response to binding as perturbations in their higher time scale dynamics.

The negatively-charged residues Glu331 and Asp332 are recognized by models utilizing vdW interactions (VDW) as well as those utilizing the higher time-scale dynamics (DYN-25 ns and DYN-100 ns). It has been shown that succinimide cyclation of Asp332 side chain alters peptide binding in PDZ3 (17). Although this residue does not interact with the binding ligand directly, it was proposed that this residue affects the local conformation of the loop, thus the electrostatic interaction between the neighboring Glu331 and the ligand is disrupted (17). Our results suggest that the Asp332 senses the ligand binding as a change in its short-range nonbonding interactions and high time scale dynamics. It is reasonably expected that the change in such properties of this residue would affect binding as well. The same paper reports that mutation of Asp332 to proline, which clearly has different flexibility and nonbonding interaction properties, affects ligand binding in PDZ3 (17).

Dynamics of Phe340 at different time scales, especially at 100 ns is shown to be affected by ligand binding according to our results. Phe340 is one of the residues that gave the highest fluctuation response upon perturbation by Perturbation Scanning Response (PRS) analysis (14). Being highlighted in PRS experiments means that the random forces put on Phe340 cause response in all protein residues. According to our results, it can be expected that limiting the dynamics of this residue might also have an allosteric effect on ligand binding.

Another PDZ3 residue that has been shown to have a relatively high response in PRS analysis in the same study is Val362 (14). According to that study, this residue is one of the highly-weighted residues in the allosteric pathway of PDZ3. This residue has a high impact on distinguishing the bound and unbound protein conformations when we incorporate its fluctuations (FLUCT) or its dynamics at 1 ns time scale (DYN-1 ns). Also, its position is important in prediction of the binding state, especially when its side chain is taken into account (GEOM). This implies that its side chain is probably more affected upon ligand binding than its backbone. Another residue in this pathway is Val386 for which the dynamics at 25 ns time scale (Dyn-25 ns) is important according to our results. These two valine residues have close side chain interactions although they are far apart in the sequence of the protein. On the other hand, Val386 interacts with Leu323, which is part of the peptide binding site interacting with peptide’s carboxylic end. Ala375, another residue close to this region, is also one of the top ten determinants in distinguishing the bound and unbound states with its low time scale (DYN-10 ps) dynamics. This residue has shown relatively high sensitivity to mutations in previous studies (15). Located at the same helix, Ala378 shows significantly high correlations in the surface area (SURF) upon binding, implying that its exposure to the solvent is changed when the peptide ligand binds to the protein. This residue interacts with Val362 (discussed above), but to our knowledge it has not been reported to be significant in any previous studies. Similarly, Ala382, which interacts with both Val362 and Val386, shows significance in models based on position (C_α_), fluctuations (FLUCT), vdW interactions (VDW), surface area (SURF) and dynamics models at all time scales. In fact, Ala382 seems to be part of a network of hydrophobic residues partnered with Ala375, Ala378, Val362 and Val386 whose dynamics at different time scales are affected upon binding (Figure 5C).

We used the top ten residues recognized by random forest models to retrain neural networks, and compared with the networks that used all non-pocket residues. As a control, we also trained the same networks with ten randomly selected residues that did not contain any of the top 10 residues nor any of the residues in the binding site. The results show that top ten residues in each model predict the binding state significantly better than the ten randomly selected residues. However, for many attributes, using the whole dataset results in significantly better predictions than using only top ten residues (Supporting Figure SI-1).

## Discussion and Conclusion

It is well known and well described that some protein residues that are located far from a binding site are affected upon ligand binding, and also, changes in conformation of these residues upon binding to allosteric ligands can potentially regulate binding affinity of the protein with its major ligands. To gain more insight into these distal allosteric residues, it is important to know whether allosteric residues communicate with the binding site in the same way or different ways. For any given allosteric protein, if all allosteric residues communicate with the binding site through same main interactions (residue attributes in this study), one just needs to focus on the location of allosteric residues when designing allosteric ligand. On the other hand, if all allosteric residues communicate with the binding site through different interactions, one should expect more complex behaviors in allosteric sites and will need to take more rationale and systematic strategies in designing allosteric ligands because different properties of different allosteric sites might be needed to induce desired changes to regulate the protein binding to its ligand. Therefore, we aimed to use hybrid models that benefit from advantages of both MD and machine learning methods to reveal how different properties of potential allosteric residues are related to ligand binding in PDZ3 domain of PSD-95 as a model protein.

The neural networks trained with different properties of MD snapshots of ligand bound and apo states trajectories show that different characteristics that describe protein residues have different potencies for distinguishing the bound and apo states snapshots. Among tested attributes, position, fluctuations and long time scale dynamics of residues are more efficient in distinguishing the binding status in PDZ3. Having various potencies in predicting the binding state of PDZ3 shows that various residue properties are affected to different extents upon ligand binding. Therefore, we can describe different properties of residues as different layers of information about the protein when it is bound to a ligand or an effector. Each of these layers has a unique contribution to predict the binding status of the protein, which implies the contribution of the residue attribute related to that layer to facilitate the ligand binding, or being affected upon binding. It is reasonable to assume that the information hidden in position, surface area, and nonbonding interactions layers might be inherited from the subtle differences in the initial structures, because the simulations were initiated from the bound and unbound crystal structures independently. However, our study shows that the dynamics and fluctuation layers of information extracted from the molecular dynamics simulations are more representative of the differences induced by ligand binding during the simulations.

Using the random forest predictive models, we represent residue-specific allostery using residue perturbation maps. Based on these maps, we demonstrate that all residues can be considered allosteric and each of them “senses” the bound ligand in a unique way. Some residues are affected in many different ways, while some are affected in a certain way. Some residues undergo changes in their nonbonding interactions with other residues; some undergo changes in their dynamics properties, while many residues are affected in a combination of these properties. Also, some residues are affected significantly, and are likely the residues being identified as allosteric residues in experimental studies, while many residues are affected only slightly. This can be considered as analogous to a social stress that results in different behavioral and emotional reactions in individuals living in the society. As occurs in a social stress, some reactions to the stress of ligand binding in a protein are more common and descriptive of the event, while some are less common. Among the common responses to the binding event in PDZ3 are perturbation of residues’ fluctuations and long time scale dynamics, and among the less common responses is the change in short time scale dynamics, which is reasonably more likely to be experienced by residues close to the binding site.

The remaining question would be why different responses are observed among protein residues upon ligand binding to the protein? A hypothetical answer would interpret unique residue response based on the unique position in sequence, space and chemical properties of the residue. Residues closer to the binding site are more likely to be affected through short-range nonbonding interactions directly from the ligand. The cascade of perturbation in such short-range interactions might fade with increasing distance from the binding site. But polar and charged residues in far distances from the binding site can be directly affected by the ligand through the electrostatic forces initiated by polar and charged moieties of the ligand. This would result in consequent perturbation in position and dynamics of these ligands, which can then start their own cascade of perturbations in their neighboring residues in time and space. If future experiments support this view about the allosteric effects in proteins, one might expect that in addition to their sequence and conformation, proteins can potentially be described with “residue perturbation maps” such as the one seen in Figure 4A, which represent the extent of specific perturbations in residue attributes. Supposing that every ligand is unique in its own chemical properties, this description of proteins would also be ligand-specific, and residue perturbation maps can also be considered as fingerprints that uniquely describe a protein when it interacts with a specific ligand. We expect that residue perturbation maps would give researchers very useful guidance when they design drugs that target allosteric binding sites. These maps provide useful information about properties of the allosteric binding sites of proteins to gain the most change of the binding affinity with their major ligands.

In this work, we chose a well-studied allosteric ligand-protein system to test our hypothesis and evaluate the feasibility of our novel approach in analyzing protein residue characteristics related to an allosteric perturbation, which is the ligand binding in this case. The results obtained with this model system show that combining machine learning models with atomistic descriptive models is a promising approach in computational structural biology, and could give us new insight into the hidden structure-activity relationships in biological macromolecules. Such combinatorial approaches inherit the benefits of each developing component, and are anticipated to lead to new important applications for these models. They can be used in future to analyze the effects of ligand binding, mutation and post-translational modifications in macromolecules.

## Acknowledgement

We acknowledge the High Performance Computing Center at the University of North Texas and the Texas Advanced Computing Center (TAAC) for providing computational resources for MD simulations and training of Deep Neural Networks related to this research work.

## Supporting Information Legends

S1. Supporting Information. Supporting Figure SI-1, details of the MD simulations and analysis, and sample analysis scripts

S2. MD input files. MD simulations parameter and input files.

S3. MD representative snapshots.

